# A Temporal Transcriptional Map of Human Natural Killer Cell Differentiation

**DOI:** 10.1101/630657

**Authors:** Aline Pfefferle, Herman Netskar, Eivind Heggernes Ask, Susanne Lorenz, Jodie P. Goodridge, Ebba Sohlberg, Trevor Clancy, Karl-Johan Malmberg

## Abstract

Natural killer cell repertoires are functionally diversified as a result of differentiation, homeostatic receptor-ligand interactions and adaptive responses to viral infections. However, the regulatory gene-circuits that define the manifold cell states and drive NK cell differentiation have not been clearly resolved. Here, we performed single-cell RNA sequencing of 26,506 cells derived from sorted phenotypically-defined human NK cell subsets to delineate a tightly coordinated differentiation process from a small population of CD56^bright^ precursors to adaptive NKG2C^+^ CD56^dim^ NK cells. RNA velocity analysis identified a clear directionality in the transition from CD56^bright^ to CD56^dim^ NK cells, which was dominated by genes involved in transcription and translation as well as acquisition of NK cell effector function. Gene expression trends mapped to pseudotime, defined by increasing entropy, identified three distinct transcriptional checkpoints, reflecting important changes in regulatory gene-circuits. The CD56^bright^ NK cell population dominated pseudotime with two distinct checkpoints separating precursors from intermediate states that gradually took on transcriptional signatures similar to CD56^dim^ NK cells. The final checkpoint occurred during late terminal differentiation of CD56^dim^ NK cells and was associated with unique divergent gene-expression trends. Furthermore, we utilized this single-cell RNA sequencing resource to decipher the regulation of genes involved in lysosomal biogenesis and found a coordinated gradual increase in the *RAB4* and *BLOC1S* gene families with differentiation into CD56^dim^ NK cells. These results identify important gene programs driving functional diversification and specialization during NK cell differentiation and hold potential to guide new strategies for NK cell-based cancer immunotherapy.

## Introduction

Natural killer (NK) cells are innate lymphocytes that play a vital role in the immune response through their ability to directly kill transformed and virus infected cells, and by orchestrating the early phase of the adaptive immune response^1^. NK cells are commonly divided into two distinct subsets based on their level of CD56 expression, eg. CD56^bright^ and CD56^dim^ NK cells, with distinct functional properties. CD56^bright^ NK cells, exhibiting an immunoregulatory role, are highly responsive to cytokine stimulation, primarily located within secondary lymphoid organs and have poor cytotoxic potential^2–4^. General consensus based on phenotypic profiling and functional and transcriptional studies identifies this NK cell population as an immature precursor to CD56^dim^ NK cells^5–9^. CD56^dim^ NK cells, making up ~90% of all circulating NK cells, express CD16 and exhibit higher cytotoxic potential coordinated through receptor-mediated input^2,10^. However, this is an oversimplified view of the NK cell repertoire. Mass cytometry profiling of NK cell repertoires at the single cell level has revealed an extensive phenotypic diversity comprising up to 100,000 unique subsets in healthy individuals^11^. Much of this diversity is based on combinatorial expression of stochastically expressed germline encoded activating and inhibitory receptors that bind to MHC and tune NK cell function in a process termed NK cell education^10,12^. Another layer of diversity reflects the continuous differentiation through several well-defined intermediate phenotypes^13^ from the naïve CD56^bright^ NK cells to the terminally differentiated, adaptive NKG2C^+^CD57^+^CD56^dim^ NK cells, associated with past infection by cytomegalovirus (CMV)^14^. Given the increasing interest to harness the cytolytic potential of NK cells in cell therapy against cancer, it is of fundamental importance to understand the molecular programs and regulatory gene circuits that drive NK cell differentiation and underlie the functional diversification of the human NK cell repertoire.

Mouse studies identified important roles for T-bet and Eomes in the differentiation from immature CD27^+^CD11b^−^ to mature CD27^−^CD11b^+^ NK cells, but did not delineate the exact intracellular signaling pathways mediating these effects^15,16^. In an attempt to characterize NK cell heterogeneity within peripheral blood and organs using single-cell RNA sequencing (scRNA-seq), Crinier et al., identified organ-specific signatures with four populations of human spleen NK cells with gene signatures along a continuum confined by the traditional CD56^bright^ and CD56^dim^ NK cell subdivision^17^. Remarkably however, only two major transcriptional subsets were found in blood-derived NK cells in both species. Through the use of bulk RNA and ChIP sequencing in human NK cells, a TCF1-LEF-MYC axis was identified in CD56^bright^ NK cells compared to CD56^dim^ NK cells where PRDM1 played a central role^18^. Furthermore, the CMV-driven adaptive NK cell responses are associated with epigenetic modifications, ultimately reflected in CD8 T cell like transcriptional profiles^19–21^. These gene regulatory programs underlying the CD56^bright^ versus CD56^dim^ NK cell phenotypic classification, provide a transcriptional basis for diverse functional roles and localization of these subsets. However, it remains to be resolved how these major NK cell subsets are related to other phenotypically defined stages of NK cell differentiation. Phenotypically, NK cells are defined using a limited number of markers, but the true heterogeneity of this population at the transcriptional level is unknown. Furthermore it remains to be examined if NK cell differentiation at the transcriptional level is a linear process and if so, what transcriptional checkpoints this may entail.

Here we used scRNA-seq and a combination of new bioinformatics tools to specifically address the developmental relationship between distinct NK cell subsets and to map the gene programs associated with transitions through discrete stages of differentiation. By sequencing equal number of cells derived from five phenotypically well-defined human NK cell subsets, our data unraveled a tightly coordinated differentiation process passing through a number of transcriptional checkpoints, associated with unique gene-expression trends and changes in functional modalities. By gaining a deeper understanding of the relationship between NK cell subsets and changes in genetic programs as cells transition through phases of NK cell education it may be possible to develop new strategies to guide NK cell differentiation towards a desired functional phenotype for cell therapy.

## Results

### NK cell differentiation defined through single cell RNA-seq

To delineate the transcriptional landscape of human NK cell differentiation, we first performed conventional bulk RNA sequencing of four sorted subsets representing distinct stages of NK cell differentiation. PCA analysis identified a clear separation between CD56^bright^, conventional and adaptive CD56^dim^ NK cells, whereas the two conventional CD56^dim^ subsets (NKG2A^+^KIR^−^ and NKG2A^−^KIR^+^) were closer together in transcriptional space (Figure 1A). The four subsets were ordered counter-clockwise in accordance with a model of NK cell differentiation laid out based on proliferative responses to cytokines, which postulated that CD56^bright^ differentiate into NKG2A^+^KIR^−^CD56^dim^, NKG2A^−^KIR^+^CD56^dim^ and finally to the most mature NKG2A^−^ KIR^+^NKG2C^+^CD56^dim^ NK cells (the latter also termed adaptive). In order to address directionality in this differentiation process and resolve the transition between these phenotypically defined cell states, we performed single-cell RNA-sequencing (scRNA-seq) on bulk NK cells and five sorted NK cell subsets, ranging from CD56^bright^ NK cells to distinct subsets of CD56^dim^ NK cells, including CD57^+^NKG2C^+^ adaptive NK cells (**Supplemental Figure 1A, 2, 3**). In total, 26,506 cells were sequenced. Single cell transcriptional data from an equal number of sequenced cells from each sample was pooled and merged in one single donor-specific t-SNE plot to examine the relationship between phenotypically defined NK cell subsets across distinct stages of differentiation (**Supplemental Figure 1B**). Because of the sorting strategy we were able to study the rare CD56^bright^ NK cells and their relationship to CD56^dim^ NK cells in greater detail. Importantly, despite the use of a limited set of markers to define the five sorted subsets, they provided a complete representation of the total bulk NK cell signature as no bulk-specific cell cluster was identified (**Supplemental Figure 1B**).

**Figure 1:**
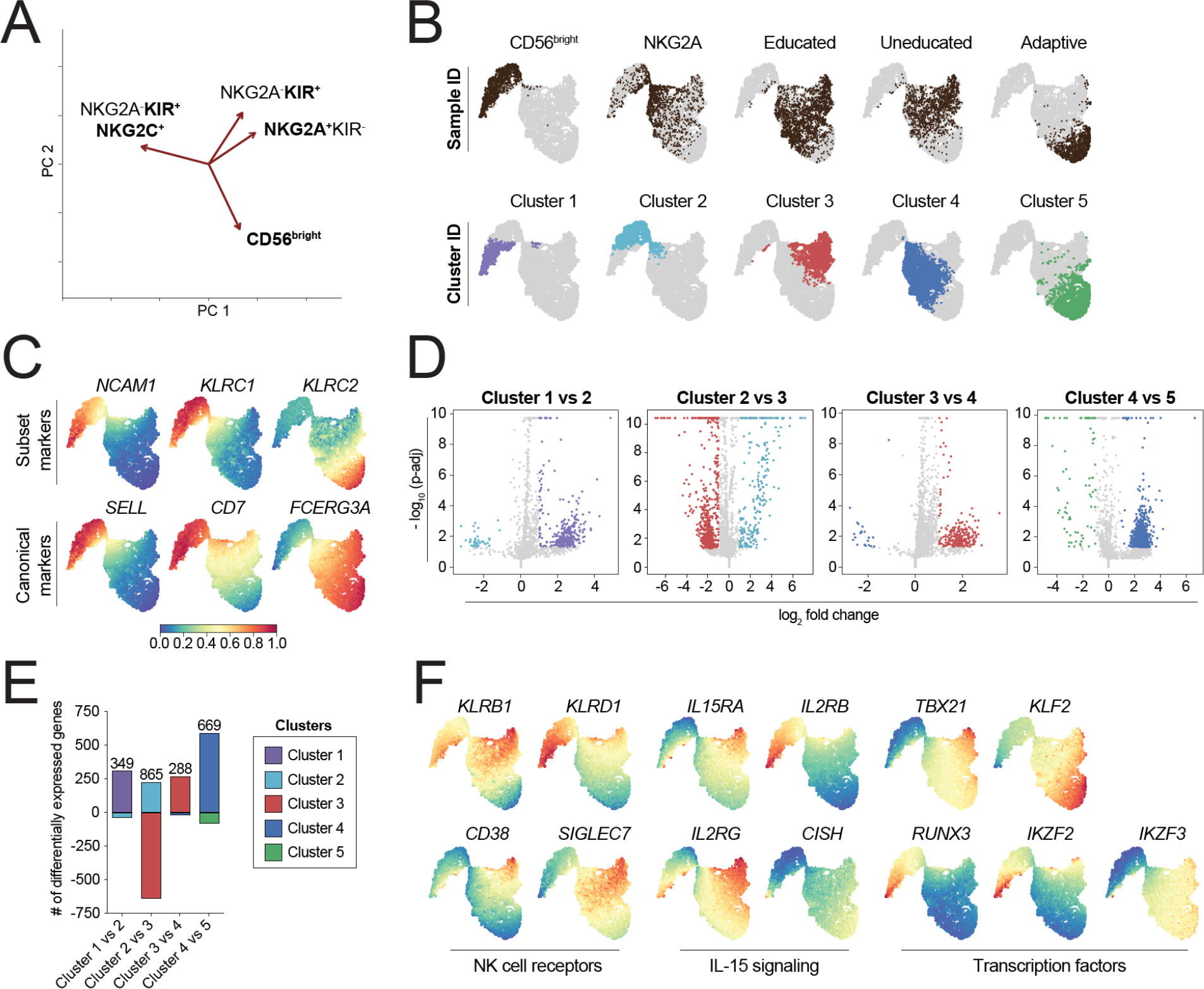
NK cell differentiation defined through bulk and single cell RNA-seq. (**A**) PCA plot of bulk RNA seq data of four discrete NK cell differentiation stages. (**B**) t-SNE plot of scRNA-seq data of five sorted NK cell subsets (CD56^bright^, NKG2A^+^ KIR^−^ CD57^−^ CD56^dim^, NKG2A^−^ self KIR^+^ CD57^−^ CD56^dim^, NKG2A^−^ non-self KIR^+^ CD57^−^ CD56^dim^, NKG2A^+^ self KIR^+^ CD57^+^ NKG2C^+^ CD56^dim^) showing transcriptional location of sorted subsets (top row) and PhenoGraph defined transcriptional clusters (bottom row). (**C**) Gene expression of selected genes displayed as a heatmap on the t-SNE plot. (**D**) Volcano plots and (**E**) summary of the number of differentially expressed genes of four comparisons between individual PhenoGraph clusters. (**F**) Gene expression of selected genes displayed as a heatmap on the t-SNE plot.

t-SNE analysis revealed two transcriptionally unique islands connected through a narrow bridge region (Figure 1B, **Supplemental Figure 4A**). Although the sorted NK cell subsets could be ordered from left to right along the previously defined maturation scheme^22^, their transcriptomes were highly overlapping with exception of the most naïve CD56^bright^ and most mature adaptive CD56^dim^ NK cell subsets. Unbiased clustering by PhenoGraph revealed two clusters in the CD56^bright^ NK cell subset and two clusters in the conventional CD56^dim^ NK cell subset (Figure 1B, **Supplemental Figure 4A**). In the first donor, adaptive NK cells represented a fifth cluster as defined by PhenoGraph that was not found in donor 2, lacking the adaptive NK cell subset. Validation of the PhenoGraph algorithm using k-means clustering yielded similar results (**Supplemental Figure 1C**). In agreement with previous data in bulk RNA-seq^23^, NK cells expressing self and non-self KIR exhibited a high degree of transcriptional overlap and together made up a significant portion of cluster 3 and 4 (**Supplemental Figure 1D**). Sorted NKG2A^+^CD56^dim^ NK cells exhibited high transcriptional variation and could be identified in all PhenoGraph-defined clusters (**Supplemental Figure 1D**). Representation of the expression of canonical differentiation markers, including *NCAM* (CD56), *KLRC2* (NKG2C), *SELL* (CD62L), *CD7* and *FCGR3A* (CD16) across the t-SNE map corroborated the gradual transitioning from naïve to more mature NK cells with progression from cluster 1 through 5 (Figure 1C).

To identify the gene signatures defining the five transcriptional clusters, we performed differential gene expression analysis between clusters defining unique steps of NK cell differentiation (Figure 1D-E). The most distal CD56^bright^ cell population, confined within cluster 1 were more transcriptionally diverse compared to cluster 2 cells and significantly increased genes in cluster 1 included *IFNG*, *OAS1*, *FGR*, *CDK6*, *CCR5* and *SLC37A1*. Visual representation of key regulatory genes also revealed high expression of *IL2RB*, *IL2RG* and *IL15RA* as well as *KLRD1*, *RUNX3* and *IKZF2* in cluster 1 (Figure 1F). The biggest transcriptional difference was observed between cluster 2 and 3, largely representing the CD56^bright^ to CD56^dim^ transition, with CD56^dim^ NK cells being more transcriptionally diverse (Figure 1D-E). Genes with significantly increased expression in cluster 2 included *XCL1*, *SELL*, *CCR1*, *LEF1*, *IL7R*, *GZMK*, *LTB*, *CD27*, *CCR7*, *MYC*, *CAPG*, *KIT*, *IL23A* and *BACH2*. Conversely, cluster 3 was defined through higher expression of *CCL3*, *CCL4*, *CCL5*, *RORA*, *GZMA*, *GZMB*, *GZMH*, *GZMM*, *FCGR3A*, *TIGIT*, *NFKBIA*, *CX3CR1*, *PRDM1*, *ZEB2*, *TFEB* and *MICB*. Cluster 3 and 4, containing a mixture of conventional, phenotypically defined, CD56^dim^ NK cell subsets, exhibited the fewest transcriptional differences (Figure 1D-E) with *CD38*, *LAIR2*, *GNAQ*, *RETSAT*, *CCDC41*, *BTRC* and *NARS2* being upregulated in cluster 3 and only *ALKBH2* in cluster 4. Overall, cluster 3 appears to represent a slightly more activated cell state within the conventional CD56^dim^ NK cell compartment with higher expression of cytokine receptors, *CD38*, *SIGLEC7*, *KLRB1*, and *TBX21*, rather than being a unique differentiation stage (Figure 1F). The comparison between cluster 4 and 5 is representative of the transition from conventional to adaptive NK cells and was characterized by a general loss of gene expression (Figure 1D-E), in line with the previously reported epigenetic reprogramming during terminal NK cell differentiation^18,19^.

Thus, analysis of scRNA-seq data from sorted NK cell subsets identified unique transcriptional clusters, which only partially overlapped with phenotypic subsets. Notably, we identified two transcriptionally-defined differentiation stages within the CD56^bright^ population and a unique cluster within the conventional CD56^dim^ population with a gene expression profile suggestive of an activated cell state.

### Continuous and coordinated transcriptional changes in pseudotime

Bulk RNA-seq of the two main NK cell populations has identified unique and evolutionary conserved regulatory programs driven by TCF1-MYC (CD56^bright^) and PRDM1-ZEB2-MAF (CD56^dim^)^18^. Plotting these regulatory genes on the transcriptional map, generated by merging single-cell transcriptional signatures of distinct NK cell subsets, suggested that the TCF-MYC axis is gradually replaced by the PRDM1-ZEB2-MAF-driven effector program (Figure 2A). Hence, we hypothesised that these genes may be used to probe directionality and temporal relationships in the differentiation process. Although sequencing of single cells only provides a snapshot in time, the ratio of spliced to unspliced mRNA content within individual cells provides the necessary data to calculate RNA velocity^24^. RNA velocity is a vector based on the time derivative of gene expression, which can predict the future state of the cell (in the range of hours) in terms of gene expression. Our dataset exhibited minimal differences in spliced and unspliced mRNA, with the exception of within the CD56^bright^ island (Figure 2B, **Supplemental Figure 4B**). Vector length, indicating the speed at which the cells are changing gene expression, increased with proximity to the bridge region linking the two transcriptional islands. Importantly, the directionality of the vector indicated a transition from the CD56^bright^ to the CD56^dim^ transcriptional island. Genes that contributed highly to the RNA velocity vector, in terms of spliced versus unspliced mRNA, included genes associated with transcription and translation (*MBNL1*, *TNRC6B*, *PARP8*, *FOXP1*, *NR3C1*), the actin cytoskeleton (*ARHGAP15*, *UTRN*, *TXK*), intracellular signaling (*PRS3*, *TNIK*), and NK cell functionality (*AOAH*, *CBLB*, *SKAP1*, *CD96*, *IL12RB2*, *CLEC2D*, *FYN, LYST*) (**Supplemental Table 1**).

**Figure 2:**
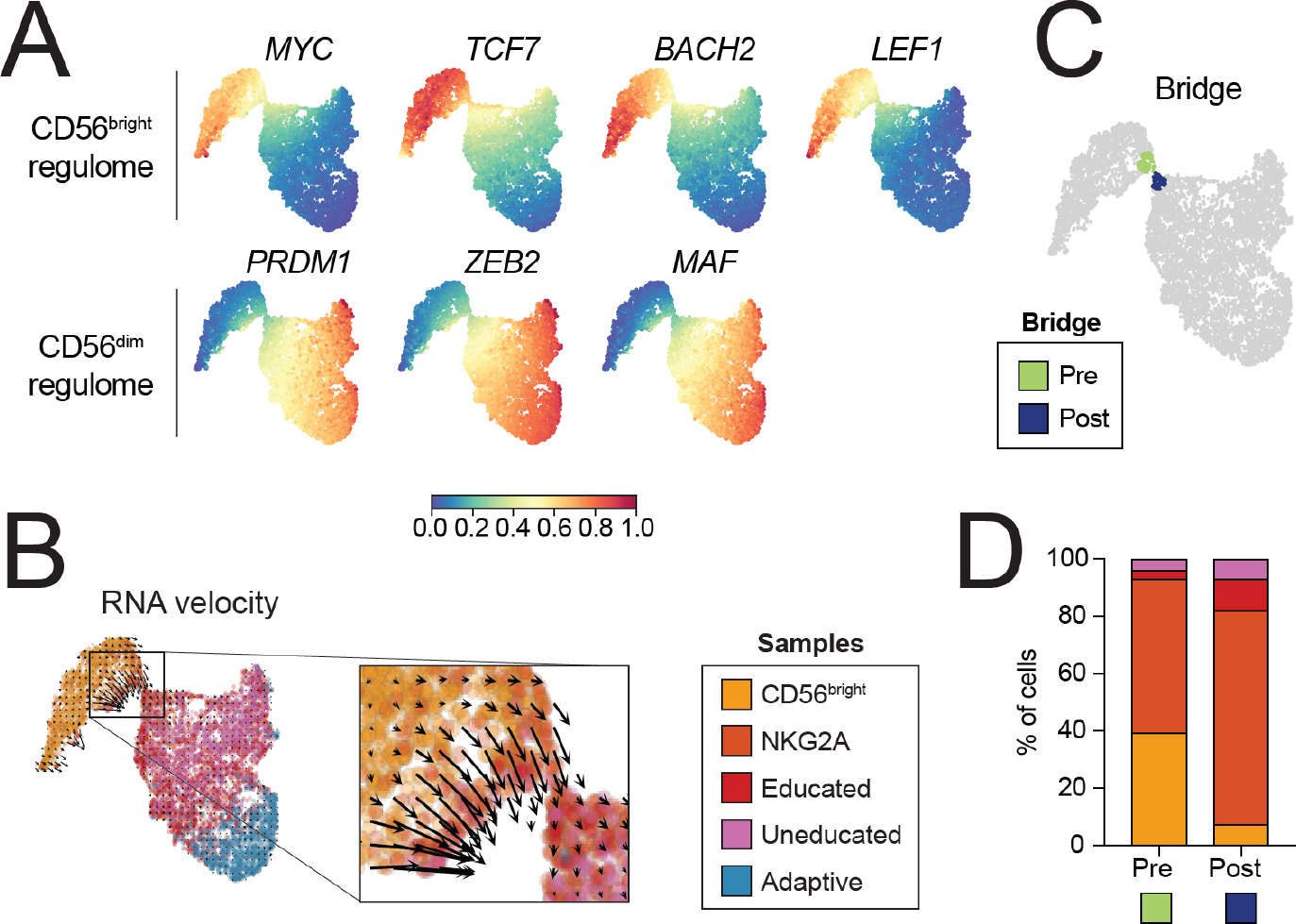
Transitioning from CD56^bright^ to CD56^dim^ NK cells. (**A**) Gene expression of selected genes displayed as a heatmap on the t-SNE plot. (**B**) t-SNE showing sample ID of sorted subsets with the RNA velocity vector overlaid and magnification of the bridge region exhibiting high RNA velocity. (**C**) Pre (green) and post (blue) bridge region clusters consisting of 100 cells each and defined based on their proximity to the bridge. (**D**) Frequency of how much each sorted subset contributes to the custom defined bridge clusters.

Careful scrutiny of the subset origin of cells localized on both sides of the bridge suggests a non-dramatic transition from CD56^bright^ to CD56^dim^ NK cells. A significant fraction of sorted NKG2A^+^CD56^dim^ NK cells were identified within the smaller predominantly CD56^bright^ island (Figure 1A, **Supplemental Figure 4A**). To further characterize the cells defining the bridge region, we identified custom clusters (pre, post) consisting of the 100 most proximal cells on each side of the bridge (Figure 2C). Despite 40% of the pre-cluster consisting of sorted CD56^bright^ NK cells, the cluster also contained 50% sorted NKG2A^+^CD56^dim^ NK cells and even a small population of KIR^+^ CD56^dim^ NK cells (Figure 2D), suggesting that changes in these commonly used phenotypic markers may be partly dissociated from underlying global transcriptional changes.

Thus, NK cell differentiation from CD56^bright^ to CD56^dim^ is associated with coordinated and yet gradual changes in phenotypic surface markers that are tightly linked to reciprocal regulatory gene circuits controlled by NK cell-specific transcription factors.

### Transcriptional checkpoints and gene-expression trends during early NK cell differentiation

To establish a transcriptional timeline of NK cell differentiation, we implemented pseudotime (Palantir) analysis, providing an unbiased approach to model trajectories of differentiating cells^25^. Palantir treats cell-fate as a probabilistic process and uses entropy to measure the changing nature of cells along the differentiation trajectory. The starting cell (highest MYC expression) was chosen based on the CD56^bright^ regulome (Figure 2A, 3A, **Supplemental Figure 4C**) identified by Collins et al^18^. Pseudotime analysis using BACH2 or LEF1 to define the starting cell yielded similar results. The Palantir algorithm identified one terminal cell, located at the tip of cluster 5 furthest from the bridge, belonging to the adaptive population (Figure 3A). In the conventional donor, the terminal cell was identified within cluster 4, belonging to the mature population (**Supplemental Figure 4C**).

**Figure 3:**
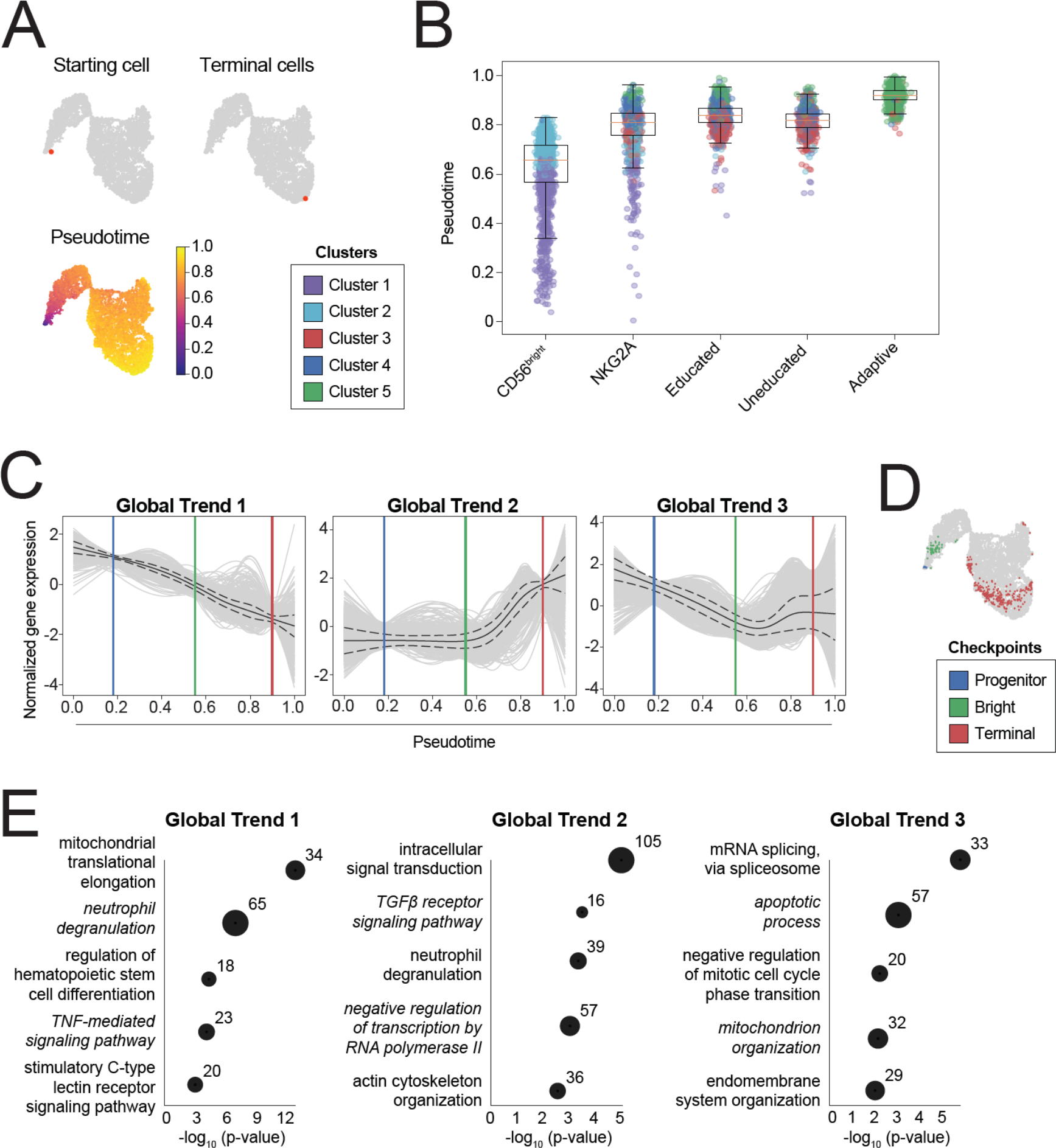
Differentiation checkpoints at distinct stages of pseudotime. (**A**) Starting cell (highest MYC expression), terminal cell and pseudotime as calculated using the Palantir algorithm. (**B**) Distribution of each sorted subset in pseudotime with colors denoting the PhenoGraph clusters each cell identifies with (box limits, upper and lower quartiles; center line, median; whiskers, quartile 1 - 1.5 * inter quartile range to quartile 3 + 1.5 * inter quartile range; points, outliers). (**C**) Global gene trends mapped onto pseudotime with colored lines denoting transcriptional checkpoints defined by low standard deviation (blue = progenitor, green = bright, red = terminal). (**D**) Visualization of cells corresponding to each checkpoint identified in pseudotime. (**E**) Gene set enrichment analysis of selected significant gene ontology (GO) terms associated with each gene trend. Significance was calculated using Fisher’s exact test followed by false discovery rate correction of the p-value. A negative log10 value of the false discovery rate adjusted p-value > 1.3 is deemed significant.

Plotting the transcriptional signatures of the sorted samples against pseudotime revealed that 75% of pseudotime, reflecting gene diversity and decreasing entropy, was occurring within the CD56^bright^ NK cell differentiation stage (Figure 3B). Distribution within the CD56^bright^ sorted cells was centered around 65% of pseudotime, with the first 60% of pseudotime corresponding to the smaller population of sorted cells grouping as cluster 1 cells (Figure 3B). This further corroborate the notion that the immature CD56^bright^ cells can be separated into two transcriptionally distinct subsets occupying distinct stages of the differentiation timeline (Figure 1A). Individual stages of CD56^dim^ differentiation were largely assigned to the final 20% of pseudotime with outliers in the NKG2A^+^CD56^dim^ NK cells being part of the earlier phase in pseudotime (Figure 3B). NK cells expressing self and non-self KIR occupied the same location within pseudotime whereas adaptive NK cells were confined to the last 10% of pseudotime (Figure 3B).

Having established a timeline for differentiation through the use of pseudotime, generalized-additive models (GAMs) were fitted on cells ordered by pseudotime to identify common gene expression. Three common global gene trends were identified, containing between 713 to 1181 genes (Figure 3C). Distinct checkpoints (blue, green, red line) with low gene expression diversity were identified in all gene trends at the same timepoints, indicative of differentiation checkpoints (Figure 3C). In addition to low gene expression diversity, these transcriptional checkpoints identified the timepoints where the gene expression changed directionality. When mapped back to the t-SNE plot, the checkpoints were located at the tip of the CD56^bright^ island (blue), prior to the transition between cluster 1 and 2 (green) and at the transition into the adaptive population (red) (Figure 3D). Hence major transcriptional changes are occurring early after entering the NK cell lineage, during differentiation within the CD56^bright^ NK population and upon transitioning into the adaptive stage. Importantly, the checkpoint within the bright population already occurs in the latter half of pseudotime, again highlighting that a dominating part of transcriptional changes occur within the CD56^bright^ population (Figure 3C).

Gene set enrichment analysis (GSEA) was utilized to characterize the main transcriptional programs associated with each global gene trend (Figure 3E). Trend 1 included genes whose expression negatively correlated with pseudotime and which belonged to processes involved in mitochondrial translational elongation, regulation of hematopoietic stem cell differentiation, TNF-mediated signaling, and stimulatory C-type lectin receptor signaling. Genes included in trend 2 are initially stable but steadily increase from the bright checkpoint to the terminal checkpoint. Transcriptional programs associated with this expression trend included intracellular signal transduction, TGFβ receptor signaling, neutrophil degranulation, negative regulation of transcription (RNA polymerase II) and actin cytoskeleton organization. The final trend, global trend 3, contained genes whose expression decreased until the bright checkpoint and then remained relatively stable until the terminal checkpoint. GO terms associated with these genes include mRNA splicing (via spliceosome), apoptotic process, negative regulation of mitotic cell cycle phase transition, mitochondrion organization and endomembrane system organization (Figure 3E).

### Diversified gene-expression patterns during terminal NK cell differentiation

As the global pseudotime analysis was dominated by CD56^bright^ NK cells we next zoomed in on the later time-points (> 80% of pseudotime), where higher standard deviation highlighted a poorer fit of the identified global gene trends (Figure 3C). Hence, transcriptional programs within the CD56^dim^ population were to a certain degree uncoupled from transcriptional programs defining NK cell differentiation within the CD56^bright^ stage. To dissect which transcriptional programs accounted for this variation observed in the gene trends, we performed new clustering only taking pseudotime > 80% into account (Figure 4A). Three new CD56^dim^ trends (dim trends) were identified, consisting of two down-trending and one up-trending trend, in line with the general decrease in gene expression observed in the transition from conventional CD56^dim^ to adaptive cells^19–21^.

**Figure 4:**
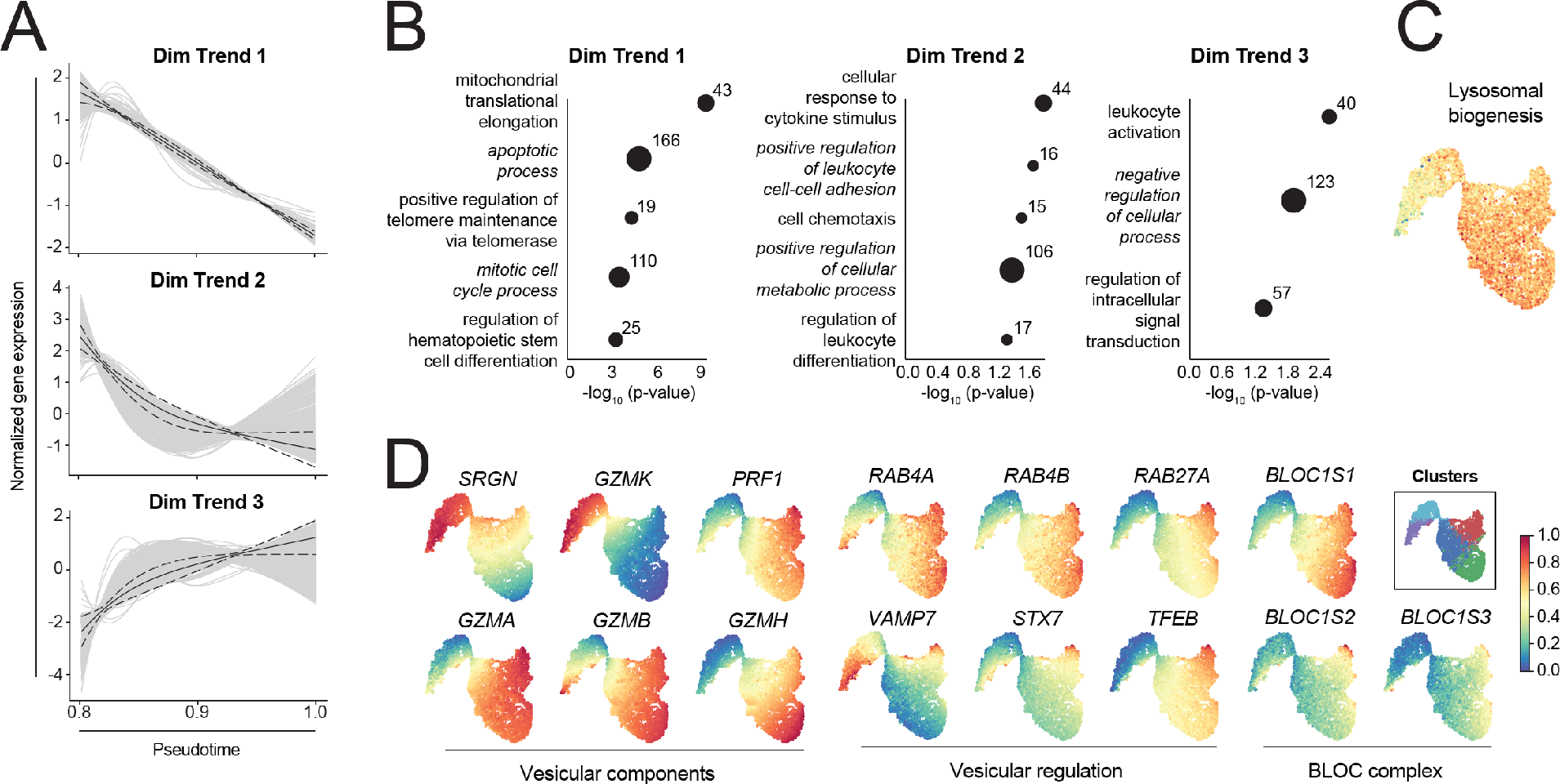
Establishment of a functional template during CD56^dim^ NK cell differentiation. (**A**)Dim gene trends mapped onto pseudotime, only taking pseudotime > 80% into account. (**B**)Gene set enrichment analysis of selected significant gene ontology (GO) terms associated with each gene trend. (**C**) Factor analysis of genes associated with lysosomal biogenesis. (**D**) Gene expression of selected genes displayed as a heatmap on the t-SNE plot. Significance was calculated using Fisher’s exact test followed by false discovery rate correction of the p-value. A negative log10 value of the false discovery rate adjusted p-value > 1.3 is deemed significant. Numbers in the plot represent number of genes identified within each GO term based on the DEGs analyzed.

Dim trend 1, the largest trend in terms of gene number (2575 genes), contained genes that steadily decreased within the later stages of pseudotime. Gene ontology terms associated with these genes include mitochondrial translation elongation, apoptotic process, positive regulation of telomere maintenance via telomerase, mitotic cell cycle process and regulation of hematopoietic stem cell differentiation (Figure 4B). Although initially decreasing, genes included in dim trend 2 were maintained at low expression from 90% of pseudotime onwards and associated with cellular response to cytokine stimulus, positive regulation of leukocyte cell-cell adhesion, cell chemotaxis, positive regulation of metabolic process and regulation of leukocyte differentiation. Lastly, dim trend 3 contained genes which steadily increased in the final stages of pseudotime, representing a clear minority. Gene ontology terms associated with these genes include leukocyte activation, negative regulation of cellular process and regulation of intracellular signal transduction.

We recently identified a role for lysosomal remodelling in the functional tuning of human NK cells during education^23^. Self-KIR^+^ NK cells show a non-transcriptional accumulation of granzyme B stored in large dense-core secretory lysosomes, which are released upon target cell recognition^23^. However, such dynamic changes to the lysosomal compartment with consequences on the ability to store granzyme B and perforin, requires a continuous and transcriptionally-regulated biogenesis of lysosomes and effector molecules as well as tight coordination of genes involved in synapse formation and degranulation. Indeed, whereas the dominating gene expression trends within the CD56^dim^ stages of differentiation (> 80% of pseudotime) show a gradual decrease in gene expression, genes involved in lysosomal biogenesis and effector function largely followed dim trend 3, which increases with differentiation and thereby provide a template for further functional tuning during NK cell education. We performed factor analysis on genes involved in lysosomal biogenesis and projected these onto the temporal transcriptional map of human NK cell differentiation (Figure 4C). Factor analysis revealed lowest expression in cluster 1, whereas the CD56^dim^ clusters (cluster 3-5) exhibited the highest expression. Next, we zoomed in on individual genes contained within the factor analysis which are important vesicular components and involved in their regulation and biogenesis (BLOC complex) (Figure 4D). The IL-15 inducible genes *PRF1* (perforin), *GZMA* (granzyme A), and *GZMB* (granzyme B) were highly expressed in cluster 3, in line with higher *IL15RA* and *IL2RG* expression in this cluster (Figure 1E). *GZMA*, *GZMB* and *GZMH* (granzyme H) were also highly expressed within cluster 5. *SRGN* (serglycin) and *GZMK* (granzyme K) exhibited a reverse gene expression pattern, being highly expressed within cluster 1 and 2. The Rab proteins (*RAB4A*, *RAB4B*, *RAB27A*) play a role in vesicular and protein trafficking as well as granule exocytosis, maturation and docking at the immune synapse. These genes exhibited higher expression within the CD56^dim^ island, with some outliers within cluster 1 also exhibiting higher expression. *VAMP7* and *STX7* are important for cytotoxic granule exocytosis in NK cells and were highly expressed within cluster 1 and 3, again in line with increased IL-15 signaling in these cells. Lysosomal exocytosis is regulated through calcium signaling and at the transcriptional level by *TFEB,* which is further regulated through phosphorylation^26^. TFEB expression greatly increased after transitioning into the CD56^dim^ island, with highest expression found in cluster 3. Lastly, the BLOC complex, consisting of *BLOC1S1*, *BLOC1S2* and *BLOC1S3*, was also higher expressed within the CD56^dim^ island. This complex is important for normal biogenesis of lysosome-related organelles, such as granules and for their intracellular trafficking.

## Discussion

We report a compact description of the transcriptional diversification at the single cell level during human NK cell differentiation. By enriching for less frequent, but phenotypically well-defined NK cell subsets, we could elucidate key regulatory gene programs within both the CD56^bright^ and the CD56^dim^ subset, as well as the developmental relationship of intermediate cell states. Pseudotime analysis highlighted the dominant role CD56^bright^ NK cells play during NK cell differentiation, with two out of three transcriptional checkpoints occurring within this small population of cells. In line with previous reports on CMV-driven epigenetic reprogramming of terminally differentiated adaptive NK cells^19–21^, the transition to this stage represented a major transcriptional checkpoint during NK cell differentiation.

The view that NK cells, like T cells, undergo a continuous process of NK cell differentiation is recent and mostly based on phenotypic and functional classification of discrete subsets^5^. Most evidence suggest that the CD56^bright^ NK cell subset is the most naïve and gives rise to the more mature CD56^dim^ NK cells which may then undergo further differentiation towards more terminal stages, a process that is accelerated by CMV infection^27^. Numerous studies have been performed in mice lacking NK specific transcription factors, using lineage tracing in macaques and in humans with immunodeficiencies, but the transcriptional identities and relationships between the manifold of putative intermediate cell states of NK cell differentiation remains elusive^5–9^.

The CD56^bright^ NK cell subset has a unique functional phenotype and tissue localization. Although infrequent in peripheral blood, they make up the large majority of NK cells within secondary lymphoid organs^4^. In line with previous reports, t-SNE analysis of a comprehensive scRNA-seq dataset from multiple sorted NK cell subsets identified two main transcriptional islands of NK cells^17^. Whereas all CD56^bright^ NK cells were confined to the smaller island, a small but definite collection of sorted CD56^dim^ NK cells, primarily NKG2A^+^KIR^−^CD57^−^, grouped transcriptionally within the CD56^bright^ population. While we cannot rule out the possibility of small numbers of CD56^bright^ NK cells contaminating our sorted population, their low frequency within the total NK cell population cannot account for the near 20% of CD56^dim^ NK cells observed within cluster 2. We have previously observed transcriptional reprogramming of educated NK cells to a more immature transcriptional signature in response to IL-15 induced proliferation which also correlated with NKG2A expression^28^. Hence, NKG2A^+^CD56^dim^ NK cells exhibiting a CD56^bright^ transcriptional signature could represent cells having undergone transcriptional reprogramming. Alternatively, these cells could represent CD56^bright^ NK cells that have downregulated CD56 expression at the surface level prior to further transcriptional changes occurring. CD56 is an adhesion molecule and has been associated with formation of a developmental synapse and distinct migratory behavior depending on the density of CD56 on the cell’s surface^29^. Hence, downregulation of CD56 surface expression could result in altered receptor input as a result of synapse formation, ultimately leading to the acquisition of a CD56^dim^ transcriptional signature. Although the bridge between transcriptional islands marked a clear decrease in *NCAM1* (CD56) expression, a gradual decreased was already observed prior to the bridge occurring within the late CD56^bright^ NK cell population. *FCERG3A* (CD16) is normally expressed on CD56^dim^ cells but can also be used to define a population of functional intermediate CD56^bright^ cells^30^. Although CD16 was not included in the sorting panel, *FCERG3A* is only significantly differentially expressed between cluster 2 and 3, despite a moderate decrease from cluster 1 to 2. Hence this functionally intermediate CD16^+^CD56^bright^ NK cell population may contribute but does not solely define the two CD56^bright^ clusters identified here.

RNA velocity, a recently described approach to predict future cell states^24^, confirmed a transcriptional transition from CD56^bright^ into CD56^dim^ NK cells occurring via transition over the bridge separating the two transcriptional islands. Among the top genes contributing to the RNA velocity vector were genes associated with NK cell receptor signaling and functionality (*LYST*, *CBLB*, *CD96*, *CLEC2D*, *SKAP1, FYN*). *LYST* is a regulator of lysosomal trafficking, and mutations in this gene leads to a lysosomal storage disorder, Chediak-Higashi syndrome, characterized by defects in NK cell degranulation leading to the accumulation of large granules within these patients^31,32^. *CBLB*, has been linked to inhibitory NK cell signaling through its modulation of LAT, a substrate of tyrosine phosphatase SHP-1^33^. Hence, Cbl ubiquitin ligase encoded for by *CBLB* is important for mediating inhibitory receptor input which is essential for NK cell functionality. Another inhibitory receptor on NK cells is CD96, which along with TIGIT and DNAM-1 can bind PVR (CD155). CD96 is expressed at the protein level upon NK cell activation^34^ and competes with DNAM-1 (CD226) for binding to their common ligand, with CD96 exhibiting an inhibitory function leading to decreased cytokine release^35^. LLT1 (*CLEC2D*) is a C-type lectin that functions as the ligand for CD161 (*KLRB1*). CD161 expression on NK cells has been linked to cytokine-responsiveness and blocking of LLT1 enhanced cytotoxicity against breast cancer cells^36,37^. In T cells, the adaptor protein SKAP-55 (*SKAP1*) links the T cell receptor to signaling via LFA-1 which is also expressed on NK cells. Furthermore, *SKAP1* can also form homodimers and could play a similar role in NK cells, where LFA-1 signaling has been linked to education^38,39^. Lastly, *FYN* is a Src kinase involved in signaling through PI3K and ERK1/2, leading to increased cytotoxicity via polarization of perforin in NK cells^40^. Transitioning across the bridge region was a gradual transition phenotypically, with defining transcriptional changes, as identified by RNA velocity occurring just prior to this region. In particular, these transcriptional changes occurred in genes having major functional implications for NK cell cytotoxicity.

A surprising finding was the dominance of the CD56^bright^ population in pseudotime and the two transcriptionally unique clusters. The first 60% of pseudotime consisted of cluster 1 cells and two of the three major transcriptional changes (checkpoints) occurred within this population and as it transitioned into cluster 2 cells. This highlights the transcriptional diversity within the CD56^bright^ population and identifies the group of cells at the beginning of pseudotime as transcriptionally unique. This small population was characterized by high *KLRB1* (CD161) *KLRD1* (CD94), *NCAM1* (CD56), *CD7*, IL15RA (CD215), *IL2RB* (CD122), *IL2RG* (CD132), *RUNX3* and *IKZF2* (Ikaros) expression. *IKZF2* has been shown to be important for NK cell precursors within the liver^41^. Rather surprising, cells in cluster 1 at the beginning of pseudotime exhibited relatively high levels of *GZMB* comparable to those in CD56^dim^ NK cells. In mice, Granzyme B mRNA is abundant in NK cells, but only translated upon cytokine stimulation^42^. In humans, CD56^dim^ NK cells express high amounts of Granzyme B compared to CD56^bright^ cells, but both can increase their expression levels in response to cytokine stimulation^43,44^. High *GZMB* expression in this early CD56^bright^ precursor state could be due to high cytokine responsiveness, as these cells also expressed high amounts of *IL2RB* and *IL2RG*. NK cell development is also dependent on cytokine priming, whereby *IL2RB* and *IL15RA* knockout mice were deficient of NK cells^45,46^. To be able to eliminate false-negatives (drop-outs) in our dataset, a result of technical limitations of scRNA-seq combined with low RNA content in resting NK cells, we implemented MAGIC to help visualize gene expression across the t-SNE map. Notably, MAGIC was not used for any other downstream analysis, effectively avoiding the generation of false positives which can be introduced by such imputation tools^47^.

The second transcriptional checkpoint was less defined in terms of standard deviation but coincided with a change in gene expression within the trends analyzed. This correlated with transitioning between cluster 1 and 2 within the CD56^bright^ NK cell population and was characterized by a decrease in *KLRB1* (CD161), *KLRD1* (CD94), *IL15RA* (CD215), *IL2RB* (CD122), *IL2RG* (CD132), *RUNX3*, *IKZF2* (Ikaros), *MYC* and *LEF1*. The decrease in cytokine receptors combined with *RUNX3* indicates reduced sensitivity to cytokine signaling, which is further reduced upon the CD56^dim^ transition^48^. This is in line with *CISH* (CIS), a negative regulator of IL-15 signaling, having the lowest expression in cluster 2^49^. Expression of effector molecule genes, including *PRF1* (perforin), *GZMA* (granzyme A), *GZMB* (granzyme B), *GZMH* (granzyme H) were also reduced in cluster 2. Overall, transitioning from cluster 1 to 2 was accompanied by a decrease in transcriptional heterogeneity, resulting in a narrower transcriptional profile in these later stages CD56^bright^ cells.

Although the conventional CD56^dim^ population exhibits a high degree of heterogeneity, both phenotypically and functionally, it only consisted of two transcriptionally defined clusters, clusters 3 and 4. These two clusters had a similar distribution, mainly containing KIR^+^ and a small proportion of NKG2A^+^CD56^dim^ NK cells. Both clusters exhibited higher expression of *IKZF3* (Aiolos) and *TBX21* (T-bet) compared to CD56^bright^ clusters, important transcription factors for maturation of NK cells in the periphery^15,50,51^. When compared to cluster 4, cluster 3 exhibited increased gene expression which was indicative of an activated genotype, characterized by high *SIGLEC7*, *CD38*, *IL15RA* (CD215), *IL2RB* (CD122) and *PRF1* (Perforin) expression, rather than a separate stage of differentiation^52^. NK cells express IL-15RA at detectable levels on the surface after IL-15 stimulation^53^. Similarly, CD38 and Perforin are both upregulated upon IL-15 stimulation. Cluster 3 cells therefore appears to represent activated cells, which are responsive to cytokine stimulation, in line with them occupying a slightly earlier pseudotime compared to cluster 4. In agreement with bulk RNA-seq in both mouse and human NK cells^23,54^, we observed no unique transcriptional signatures between self KIR (educated) and non-self KIR (uneducated) NK cells and both sorted populations occupied the same space in pseudotime^23^. Remodelling of the lysosomal compartment has been shown to be important for the increased functionality observed in educated NK cells^23^. Here, we observed an increase in lysosomal biogenesis in the later stages of pseudotime, in line with increased functionality within the CD56^dim^ NK cells. Thus, NK cell differentiation establishes a functional template through a tightly controlled and transcriptionally regulated increase in the expression of genes involved in lysosomal biogenesis and exocytosis alongside increased expression of effector molecules such as granzyme B and perforin. Acquisition of self or non-self KIRs during later stages of differentiation sets the cells on different functional trajectories during education, which involves a non-transcriptional remodelling of the lysosomal compartment that ultimately changes the cytotoxic payload of the cell^23^.

Adaptive NK cells clustered independently and uniquely identified within the last 10% of pseudotime. Transitioning into cluster 5 was accompanied by the third and final checkpoint, highlighting the important transcriptional changes occurring at this stage of differentiation. Compared to conventional CD56^dim^ NK cells, the global transcriptome of adaptive NK cells was highly reduced. This is in line with epigenetic silencing that has been described for this population of terminally mature NK cells^19–21^. Gene expression of *SYK*, *CD38* and *KLRB1* (CD161) was reduced while *ZEB2*, *KLF2*, *PRDM1* (BLIMP-1), *KLRC2* (NKG2C), *GZMH* (Granzyme H) were highly expressed.

Here we have applied new bioinformatic tools to a unique single-cell RNA sequencing dataset in order to identify a temporal transcriptional map of human NK cell differentiation. Mapping gene expression trends to pseudotime allowed for the identification of distinct transcriptional checkpoints highlighting important transcriptional changes during NK cell differentiation. Two previously undescribed transcriptional populations within the CD56^bright^ subset were identified and dominated the differentiation timeline. This dataset provides a valuable tool to identify important gene programs that drive functional diversification and specialisation during NK cell differentiation. Such knowledge hold potential to guide the development of new strategies for NK cell-based cancer immunotherapy.

## Materials & Methods

### Cell processing

Peripheral mononuclear cells (PBMC) were isolated using density gradient centrifugation from anonymized healthy blood donors (Oslo University Hospital) with informed consent as approved by the regional ethics committee in Norway (scRNA-seq) and Sweden (bulk RNA-seq) (2015/2095, 2016/1415-32, 2018/2485). Donor-derived PBMCs were screened for KIR education and adaptive status using flow cytometry. NK cells were purified using an AutoMACS (DepleteS program, Miltenyi Biotec) and prior to overnight resting in complete RPMI (10% Fetal calf serum, 2mM L-glutamine) at 37°C/5% CO_2_.

### Flow cytometry screening

PBMC were stained for surface antigens and viability in a 96 V-bottom plate, followed by fixation/permeabilization and intracellular staining at 4°C. The following antibodies were used in the screening panel: CD3-V500 (UCHT1), CD14-V500 (MφP9), CD19-V500 (HIB19), Granzyme B-AF700 (GB11) from Beckton Dickinson; CD57-FITC (HNK-1), CD38-BV650 (HB-7), KIR3DL1-BV421 (DX9) from BioLegend; KIR2DL1-APC-Cy7 (REA284), CD158a,h-PE-Cy7 (11PB6), from Miltenyi Biotec; CD158b1/b2,j-PE-Cy5.5 (GL183), NKG2A-APC (Z199), CD56-ECD (N901) from Beckman Coulter. LIVE/DEAD Fixable Aqua Dead Stain kit for 405 nM excitation (Life Technologies) was used to determine viability. Samples were acquired on an LSR-Fortessa equipped with a blue, red and violet laser and analyzed in FlowJo version 9 (TreeStar, Inc.).

### FACS sorting

Cells were harvested and surface stained with the following antibodies: CD57-FITC (HNK-1) from BioLegend; KIR3DL1S1-APC (Z27.3.7), CD56-ECD (N901), CD158b1/b2,j-PE-Cy5.5 (GL183), from Beckman Coulter; KIR2DL1-APC-Cy7 (REA284), NKG2C-PE (REA205), NKG2A-PE Vio770 (REA110) from Miltenyi Biotec. 12,000 cells were directly sorted into Eppendorf tubes at 4°C for each sample (**Supplemental Figure 1A**) using a FACSAriaII (Beckton Dickinson).

### Bulk RNA sequencing

Four populations (CD56^bright^, NKG2A^−^KIR^−^CD56^dim^, NKG2A^−^KIR^+^CD56^dim^, and NKG2A^−^ KIR^+^NKG2C^+^CD56^dim^) were sorted from six individual healthy blood donors. Sequencing was performed using single-cell tagged reverse transcription (STRT)^55^. Principle component analysis (PCA) was used to generate a biplot of the four sequenced subsets.

### Single-cell RNA sequencing

Following sorting, cells were kept on ice during the washing (PBS + 0.05% BSA) and counting step. 10,000 cells were resuspended in 35 μL (PBS + 0.05% BSA) and immediately processed at the Genomics Core Facility (Oslo University Hospital) using the Chromium Single Cell 3’ Library & Gel Bead Kit v2 (Chromium Controller System, 10X Genomics). The recommended 10x Genomics protocol was used to generate the sequencing libraries, which was performed on a NextSeq500 (Illumina) with 5~ % PhiX as spike-inn. Sequencing raw data were converted into fastq files by running the Illumina`s bcl2fastq v2.

### Quality control and normalization of scRNA-seq data

Data cleaning steps were first carried out whereby cells not expressing a minimum of 1000 molecules and genes expressed by less than 10 cells were filtered out. The data was normalized using log transformation based on the total expression of the gene in the sample, the default normalization method implemented in the Palantir library. Feature selection was carried out to select high cell-to-cell variance, implementing a cutoff of log2-fold change > 2, whereby only significantly differentially expressed genes were selected.

### Dimensionality reduction of scRNA-seq data

A number of dimensionality reduction methods were implemented, including principle component analysis (PCA) for initial noise reduction and non-linear diffusion maps to estimate a lower dimensional manifold that could be implemented for further downstream analysis. For visualization purposes, t-SNE and UMAP were utilized^56,57^.

### Gene expression imputation of scRNA-seq data

Markov affinity-based graph imputation of cells (MAGIC) was utilized to de-noise the data in order to optimize the gene expression analysis for visualization on the t-SNE maps^58^. The imputed data matrix was not used for further downstream analysis and was not used for computation of the differentially expressed genes (DEG).

### Differentiation trajectories and gene trend analysis of scRNA-seq data

Palantir was used to carry out the trajectory analysis and pseudotime calculations^25^. The starting cells was identified as having the highest MYC expression, for which the imputed data matrix was used to eliminate selection of outlier cells. The Palantir algorithm calculates the probability of each individual cell to end up in each of the inferred terminal states, whereby only one terminal state was identified in both donors. Generalized-additive models (GAMs) fitted on cells ordered by pseudotime were used to calculate gene trends, where the contribution of cells was weighted by their probability to end up in the given terminal state as calculated by Palantir. The gene trends indicate how gene expression levels develop over the differentiation timeline. Local gene trends were calculated by zooming in on a particular range of pseudotime (> 0.8).

### Clustering, differential gene expression and gene set enrichment analysis scRNA-seq data

The gene trends were clustered using the PhenoGraph algorithm and confirmed using k-means clustering, whereby the number of clusters identified by PhenoGraph was utilized as the k input parameter^59^. For differential gene expression (DEG) analysis, the SCDE package implementing the Bayesian approach, was utilized^60^. SCDE is optimized to deal with the single cell specific challenge of dropouts. A log-fold change of > 2 and adjusted p-value > 0.05 were deemed significant. Gene over-expression analysis (Gene Ontology, PANTHER) was used downstream of the gene trend clustering and DEG analysis to identify significant differences in biological pathways.

### RNA velocity of scRNA-seq data

RNA velocity was run directly on the output created by Cell Ranger, containing the count matrix and the abundance of spliced and unspliced versions of each transcript^24^. Using the velocyto Python library, the velocity vectors and locally average vector fields were calculated, which were the projected onto the same t-SNE or UMAP embedding that was used for visualizing other analysis.

### Factor analysis of scRNA-seq data

Factor analysis was utilized to obtain a single metric for a set of genes. The gene lists for specific biological functions were obtained from Gene Ontology terms and the f-scLVM method in the Python package slalom was then used for the factor analysis^61^.

## Data sharing statement

All sequencing data (bulk and sc) will be deposited at NCBI GEO depository and will be accessible with an accession number GEO: X or using the link https://www.ncbi.nlm.nih.gov/geo/query/X.

## Authorship and conflict-of-interest

A.P. performed the single-cell RNA sequencing experiments and the bulk RNA sequencing experiments. S. L. performed the RNA sequencing and library preparation. H.N. and T.C. performed the bioinformatic analysis. E.H.A. analyzed data. J.P.G and E.S. provided scientific input. H.N. and T.C. contributed to the writing of the paper. H.N., T.C, A.P and K-J.M. designed research. A.P. and K-J.M. wrote the manuscript. K-J.M. is a scientific advisor and consultant at Fate Therapeutics.

## Code availability

Custom code utilized for analysis is available on the GitHub repository, https://github.com/hernet/SingleFlow.

