## Supplementary data for "A Temporal Transcriptional Map of Human Natural Killer Cell Differentiation"

### Supplemental Figure 1

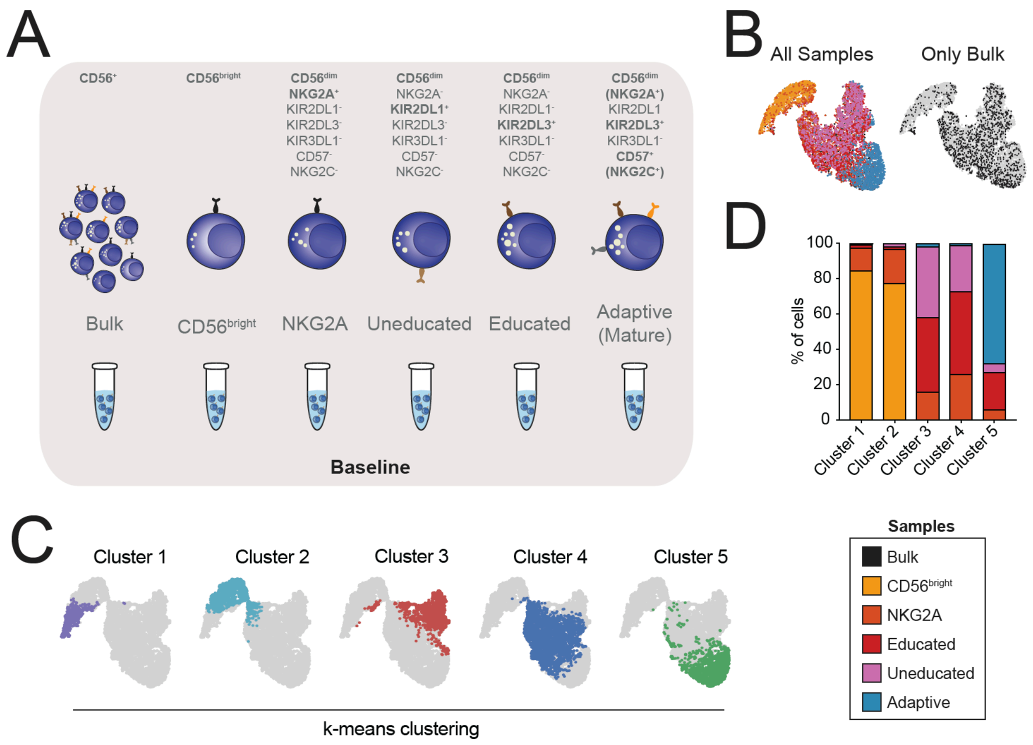


**Supplemental Figure 1: Phenotypic characterization of scRNA-seq data**

(**A**) Overview of sorted NK cell populations used for single-cell RNA sequencing including markers used for defining each population phenotypically. (**B**) t-SNE plot of all six sorted samples and only the bulk sample, with colors denoting each sample. (**C**) Validation of clusters. t-SNE plot showing transcriptional location of transcriptional clusters as defined by k-means clustering. (**D**) Frequency of how much each sorted subset contributes to the PhenoGraph defined clusters.

### Supplemental Figure 2

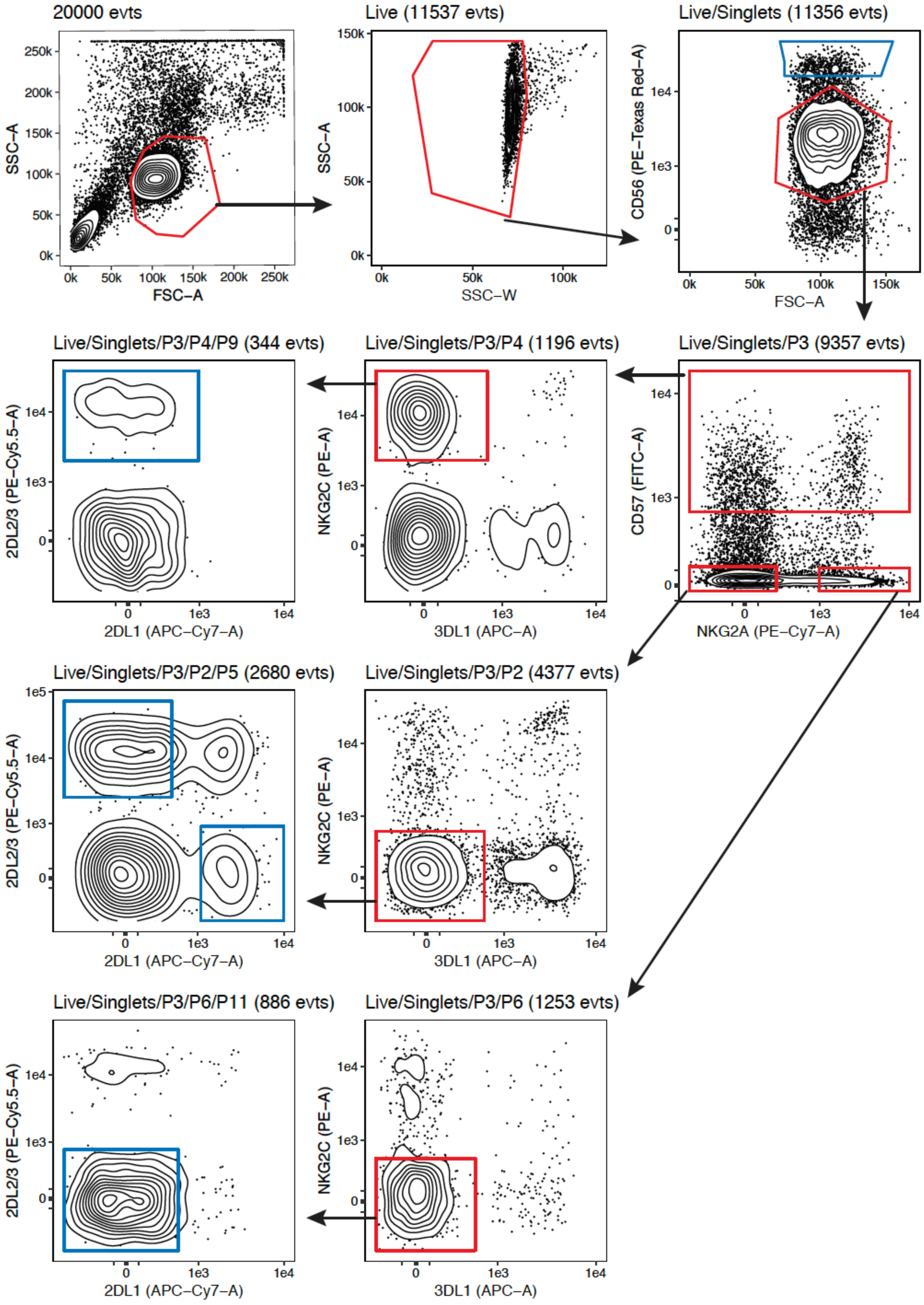

**Supplemental Figure 2: Sorting strategy in donor with adaptive NK cells**

The gating strategy used to sort five distinct NK cell populations within an adaptive NK cell donor (C1/C1). Blue gates denote sorted populations, namely $CD56^{bright}$, $NKG2A^{+}KIR^{-}CD57^{-}CD56^{dim}$, $NKG2A^{-}$self $KIR^{+}CD57^{-}CD56^{dim}$, $NKG2A^{-}$non-self $KIR^{+}CD57^{-}CD56^{dim}$, $NKG2A^{+}$self-$KIR^{+}CD57^{+}NKG2C^{+}CD56^{dim}$ NK cells.

### Supplemental Figure 3

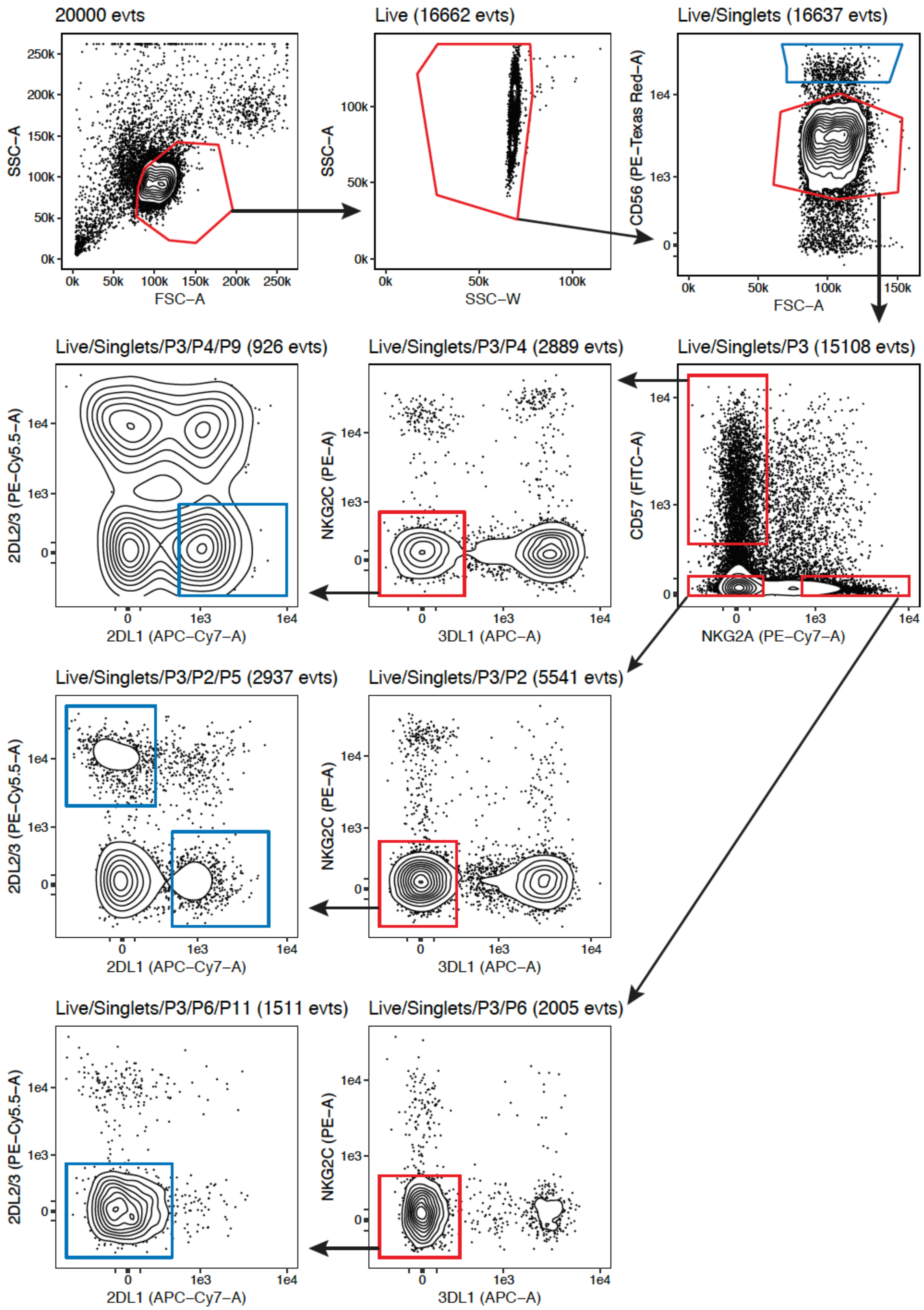

**Supplemental Figure 2: Sorting strategy in donor lacking adaptive NK cells**

The gating strategy used to sort five distinct NK cell populations within a conventional NK cell donor (C2/C2). Blue gates denote sorted populations, namely $CD56^{bright}$, $NKG2A^{+}KIR^{-}CD57^{-}CD56^{dim}$, $NKG2A^{-}$self $KIR^{+}CD57^{-}CD56^{dim}$, $NKG2A^{-}$non-self $KIR^{+}CD57^{-}CD56^{dim}$, $NKG2A^{-}$self-$KIR^{+}CD57^{+}CD56^{dim}$ NK cells.

### Supplemental Figure 4

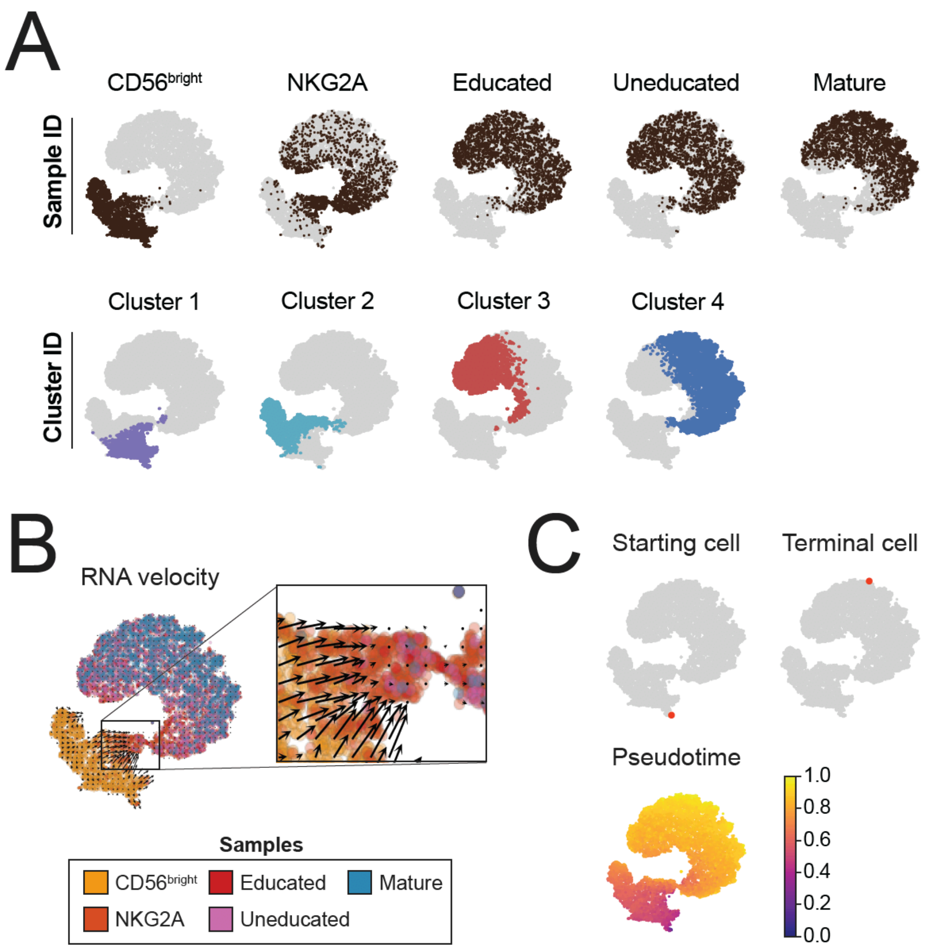


**Supplemental Figure 4: Validation of analysis in donor 2**

(**A**) t-SNE plot showing transcriptional location of sorted subsets (top row) and PhenoGraph defined transcriptional clusters (bottom row). (**B**) t-SNE showing sample ID of sorted subsets with the RNA velocity vector overlaid and magnification of the bridge region exhibiting high RNA velocity. (**C**) Starting cell, terminal cell and pseudotime as calculated using the Palantir algorithm.

| RNA velocity |
|---|
| PITPNC1 |
| MBNL1 |
| AOAH |
| LYST |
| TNRC6B |
| FOXP1 |
| CBLB |
| RPS3 |
| CLDND1 |
| SKAP1 |
| CD96 |
| UTRN |
| IL12RB2 |
| CLEC2D |
| TXK |
| FYN |
| XYLT1 |
| TNIK |
| ARHGAP15 |
| NR3C1 |
| PARP8 |

**Supplemental Table 1**. Genes highly contributing to the RNA velocity vector based on the ratio of splice versus unspliced transcripts within cells located prior to the bridge region.
